# Selective targeting of hippocampal fast-spiking basket cell interneurons via noninvasive enhancer-AAV delivery

**DOI:** 10.1101/2025.11.13.688234

**Authors:** Viktor J. Olah, Matthew J.M. Rowan

## Abstract

Accurate targeting of specific brain regions through non-invasive methods has long been a major goal of basic and translational neuroscience. Systemic delivery of AAVs expressing highly region- and cell-type-specific regulatory elements (enhancer-AAVs) continues to emerge as a tractable solution. Here we performed an independent characterization of a novel enhancer element with apparent robust transgene expression in fast-spiking basket cell interneurons in mouse hippocampus. Surprisingly, this vector did not induce expression in other brain regions harboring the same neuron class. Following intravenous administration in mice, robust labeling of FS-BCs across all hippocampal subfields was observed. We validated the FS-BC specificity in hippocampus using immunofluorescence and electrophysiological recordings. Viral labeling was confined to parvalbumin-positive cells exhibiting basket cell morphology and fast-spiking responses. This approach represents a promising avenue for both mechanistic investigation of hippocampal circuit function and potential therapeutic interventions targeting hippocampal pathophysiologies such as epilepsy, schizophrenia, and other neurological disorders in mice and potentially other species.

## Introduction

The hippocampus plays critical roles in pattern recognition, spatial navigation, and episodic memory formation, and is considered a key locus where multiple brain disorders originate. These disparate functions emerge from coordinated activity across the “trisynaptic circuit,” consisting of the dentate gyrus (DG), Cornu Ammonis 3 (CA3), and CA1 regions, where information undergoes distinct transformations at each stage (1, 2). The functional specialization of hippocampal subfields is evident in their divergent spatial coding schemes and contributions to different aspects of memory (3–5). Central to hippocampal circuit function is the precise balance between excitation and inhibition, which is orchestrated by unique inhibitory interneuron populations (6). Among these, fast spiking basket cells (FS-BC) represent a particularly important class of GABAergic interneurons that provide powerful perisomatic inhibition to principal neurons, thereby regulating network oscillations, spike timing, and thus circuit stability (7, 8). Disruption of FS-BC function has been implicated in epilepsy, schizophrenia, Alzheimer’s disease, and other neuropsychiatric disorders, underscoring the importance of understanding their specific contributions to hippocampal physiology and pathology (9–14).

Methods for investigating specific populations of hippocampal cells have improved significantly over time. Early lesion studies took advantage of the distinct anatomical organization of the hippocampal subfields (15). In contrast, more recent genetic techniques, such as using Cre mouse lines, have allowed researchers to target specific cell types (16, 17). However, Cre-based methods are limited to mouse models, often necessitate extensive crossbreeding, and cannot translate directly to clinical applications. These challenges have led to a growing interest in more versatile targeting methods, particularly enhancer adeno-associated viruses (enhancer-AAVs) (18–20). Enhancer-AAVs utilize short transcriptional regulatory elements to drive selective expression of transgenes within specific cell populations in wild-type animals (21). Notably, these elements often retain their cell-type specificity across different species, including humans, making them valuable tools for both basic research and translational applications (19, 20, 22). Despite the critical importance of fast-spiking basket cells (FS-BCs), enhancer-AAVs specifically targeting these interneurons have only recently become available (23).

Using open-source data from the Allen Institute, here we identified an enhancer-AAV with apparent specificity for FS-BCs throughout the entire hippocampus of mature mice (24). Strikingly, only minimal expression was observed in other regions within the same brains. Thus, we termed the approach *hippo BASK-IT* (hippocampal basket cell interneuron targeting). *hippo BASK-IT* provides a unique avenue for selective targeting of hippocampal basket cells using systemic AAV introduction, offering new opportunities to dissect their contributions to circuit function and their unique molecular signature (13). This approach should also accelerate evaluation of interneuron-specific cellular therapies in hippocampus. Furthermore, due to the powerful control of FS-BCs over hippocampal excitability, this enhancer-AAV is ideally suited for regulating hippocampal activity *in vivo* via chemogenetic or analogous methods.

## Results and Discussion

Using the recently developed Allen Institute brain knowledge platform (24), we identified an enhancer-AAV (AiE0476m) with a putative expression pattern reminiscent of FS-BCs which also showed remarkable specificity for the hippocampus. To test this we synthesized a new AAV plasmid incorporating two copies of the AiE0476m enhancer element sequence in tandem with additional upstream regulatory elements to boost transcriptional efficiency (18), and packaged it into the PHP.eB capsid (25) for systemic delivery. Retro-orbital injections of this vector allowed fast and easy delivery to the entire central nervous system. Approximately 4 weeks after injection (Figure 1A), mice were perfused and brains processed for imaging.

**Figure 1.**
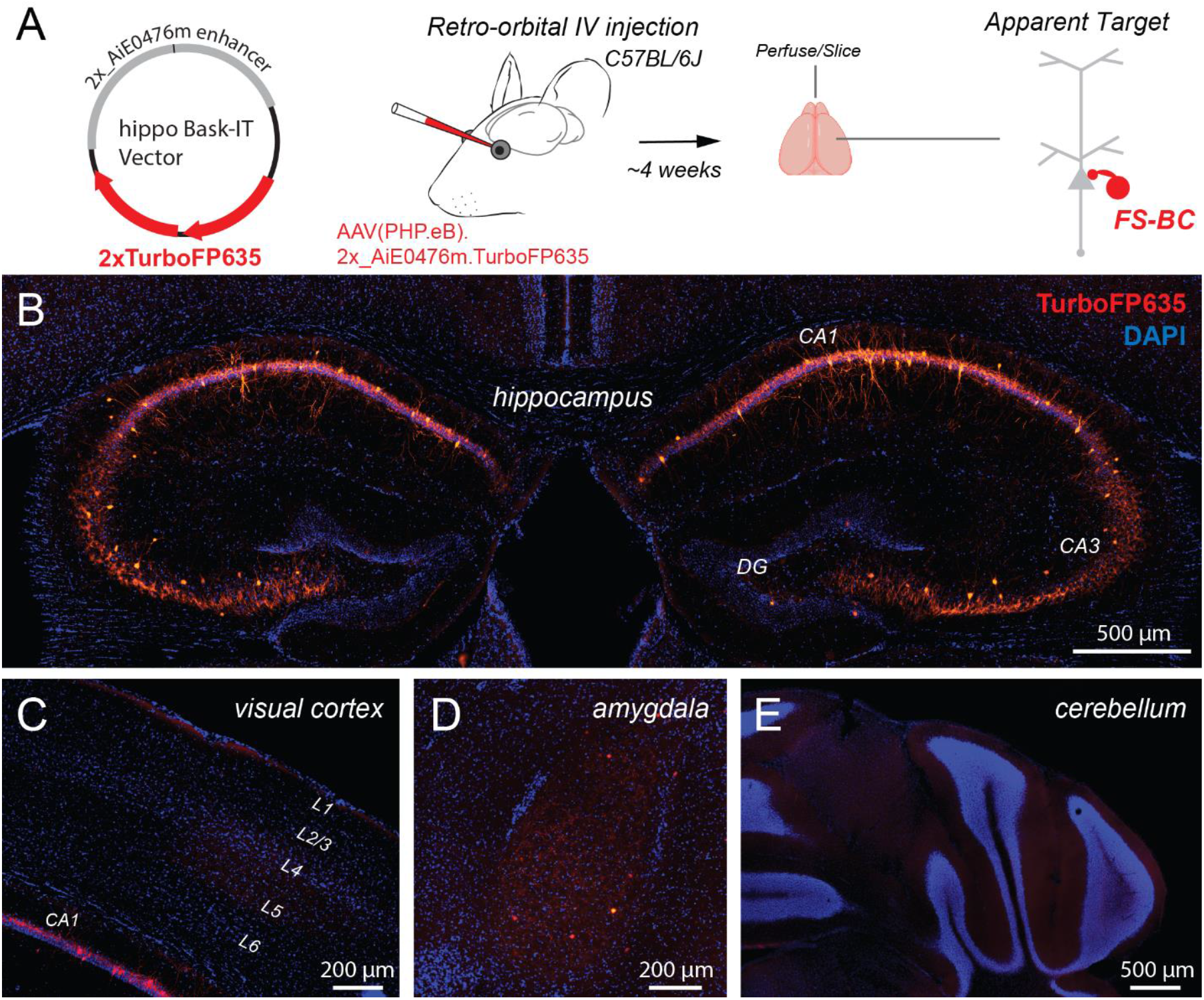
Retro-orbital injection of an enhancer-AAV results in robust hippocampal-specific labeling. **A.** Schematic illustration of the enhancer-AAV expressing the fluorophore TurboFP635 (left), the retro-orbital vector delivery method (middle), and the apparent labeling pattern in hippocampus (right). **B**. Coronal sections showing dense hippocampal labeling. **C**. Labeling (apparent lack thereof) in the visual cortex above the dorsal CA1 (note red neurons in CA1). **D**. Dim and sparse but apparent labeling in amygdala neurons. **E**. Cerebellum region showing lack of expression in the same brain.

Examination of expression throughout the brain revealed neuronal labeling in the hippocampal formation, including subfields CA1, CA2, CA3 with sparser labeling in the dentate gyrus (Figure 1B). Both the somatic position and axonal arborization of labeled neurons were highly suggestive of PV interneuron labeling. Under the same imaging conditions in the same slices, negligible labeling was observed in the cortex (Figure 2C). This was surprising, as cortical parvalbumin-expressing FS-BCs with highly analogous molecular and physiological signatures to those in hippocampus are the most numerous interneuron type in that region. This suggests a differential role for this enhancer element, and any gene expression it may regulate, in hippocampal vs cortical FS-BCs. Importantly, expression was also not observed in cerebellar neurons. This is in contrast to the recently described BiPVe3 basket cell enhancer, which drives expression more broadly including in cortex, hippocampus, and in cerebellar Purkinje cells (23). However, sparse neuronal labeling was seen in the amygdala, including the lateral and medial parts (Figure 1C,D,E).

**Figure 2.**
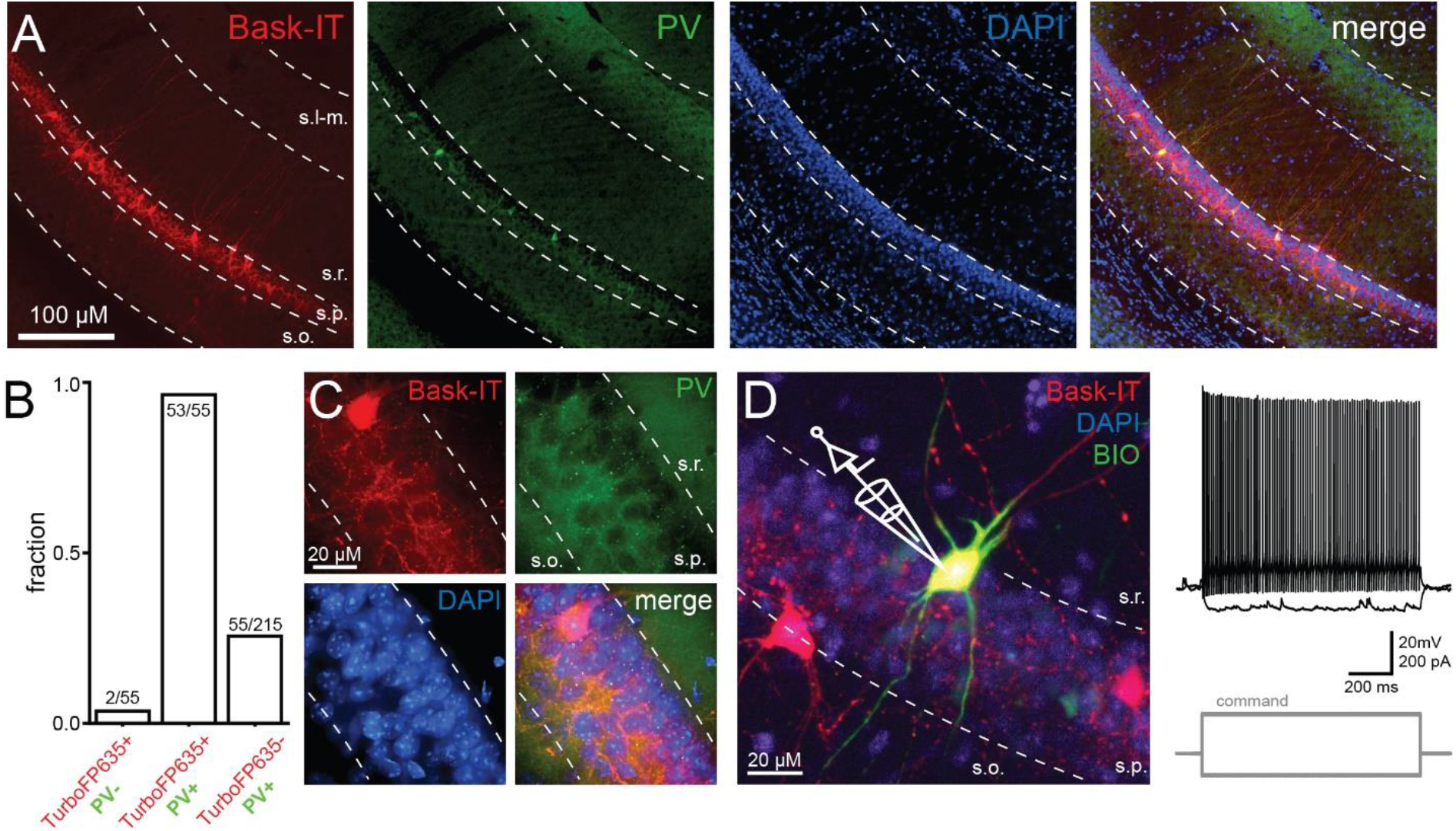
hippo BASK-IT selectively targets hippocampal fast spiking basket cells. **A.** hippo BASK-IT labeled cells are positive for parvalbumin with immunolabeling. **B**. The majority of TurboFP635+ cells are parvalbumin positive, and their number comprises 25.5% of the total hippocampal parvalbumin positive cell population. **C**. High magnification confocal images provide a detailed depiction of classical basket axons in the stratum pyramidale. **D**. Post hoc recovery of a biocytin filled TurboFP635+ cell for electrophysiology (left) and the cell’s response to hyperpolarizing and depolarizing current steps, showing fast spiking characteristics. s.o.: stratum oriens, s.p.:stratum pyramidale, s.r.: stratum radiatum, s.l-m.: stratum lacunosum-moleculare.

Closer inspection of the labelled cells in hippocampus was next performed using confocal microscopy, revealing sparse labeling of neurons located within or bordering on the stratum pyramidale in the CA3, CA2 and CA1 regions (Figure 2A) and the stratum granulosum in the dentate gyrus. The dendrites of the labeled cells were not polarized and were devoid of dendritic spines, indicating an inhibitory neuron identity. Lastly, the axonal arborization was restricted to the cell body layer, which was expected from FS-BCs (Figure 2A, C).

FS-BCs can be further identified using a combination of anatomical, neurochemical and electrophysiological markers(26). FS-BCs express the calcium buffer parvalbumin (PV)(6), which can be tested with immunohistochemistry. We found that cells expressing TurboFP635 were nearly universally positive for PV immunostaining (96.3%, n=55). TurboFP635+ cells comprised a substantial fraction of the total PV positive cell population (25.6%, n=215), indicating efficient systemic viral targeting via the PHP.eB capsid (Figure 2B). High magnification confocal imaging revealed characteristic basket axons in the stratum pyramidale surrounding DAPI labeled cell nuclei (Figure 2C). Importantly, although basket-like axonal structures could be clearly resolved, we found no indication of axonal cartridges characteristic of another major perisomatic targeting PV expressing cell population in the CA1; the axo-axonic cells(6). These axonal innervation patterns were similar in the CA3 and DG as well.

In the Cornu Ammonis regions of the hippocampus proper there are two functionally distinct cell populations that exhibit similar basket cell morphological features: the FS-BCs, and the cholecystokinin expressing basket cells (CCK-BCs)(6). Although these two cell types exhibit similar morphological features, they fulfill vastly different network functions due to their different connectivity and biophysical properties(27). FS-BCs fire at high frequencies with narrow action potentials and display minimal accommodation, while CCK-BCs have slower membrane time constants, broader action potentials and fire at low frequencies with marked frequency accommodation(28). Thus, to complement our parvalbumin immunolabeling experiments showing highly specific FS-BC targeting, we next carried out physiological characterization of TurboFP635+ neurons as well in the CA1, using fluorescent-guided patch clamp electrophysiological recordings (Figure 2D). TurboFP635+ cells showed firing behavior and AP waveform characteristics consistent with FS-BC identity (AP peak: 28.22 ± 1.98 mV, AP threshold: −47.62 ±1.35 mV, AP halfwidth: 0.27 ± 0.01 ms, AHP: 20.42 ±1.08 mV, membrane time constant: 9.81 ± 0.45 ms, maximum firing rate: 171.3 ± 16.97 Hz, n=6). Together, these results indicate that the *hippo BASK-IT* approach effectively targets FS-BCs in the hippocampus, with no observed off-target labeling. Importantly, the labeling is largely confined to the hippocampus, offering unique therapeutic and research opportunities.

GABAergic interneurons are essential for information processing. Among this functionally divergent cell population, FS-BCs are thought to have particularly important functions. This cell type represents the most abundant GABAergic cell type in the hippocampus, corresponding to ~20% percent of all inhibitory cells in this region (6), even though the number of identified hippocampal inhibitory cell types numbers more than twenty. In addition to their abundance, these cells innervate an incredibly large number of target neurons, generating large inhibitory postsynaptic conductances in their targets (29), and thereby effectively regulating a substantial portion of their surrounding network. FS-BCs have been extensively studied and are intimately linked to hippocampal rhythmogenesis, particularly to generating and maintaining gamma frequency oscillations that support cognitive processing (30–32). The importance of these interneurons is underscored by their implication in local circuit operations, learning and memory (33, 34), and sensory processing (35), with aberrant activity mechanistically linked to multiple neurological and psychiatric disorders including epilepsy, autism spectrum disorders, schizophrenia, and Alzheimer’s disease (12–14, 36–41). For example, in mice expressing Alzheimer’s pathology, parvalbumin+ basket cells in hippocampus show altered activity during sharp wave ripples, which are critical for memory consolidation (41). FS-BCs are one of the cornerstones of hippocampal physiology and pathophysiology, therefore their efficient targeting with viral methodologies is paramount for understanding neuronal circuit behaviors and for developing therapeutic avenues.

In recent decades, multiple experimental approaches have been utilized to isolate and study the hippocampus in mice, ranging from physical dissection to functional inactivation techniques. Early endeavors into understanding hippocampal function included global hippocampal lesions, which provided profound insights into the role of this brain region in behavior (42–44). More recently, experimental approaches involve microdissection of specific hippocampal subregions and strata using microsurgical techniques (45). To functionally isolate the hippocampus and overcome limitations of permanent lesions, which can impact multiple memory processes and allow time for compensatory changes in other brain regions, acute inactivation techniques have been developed using intrahippocampal injection of muscimol, a GABA receptor agonist, to temporarily silence hippocampal activity at specific stages of memory tasks (46–48). More recently, chemogenetic approaches have enabled researchers to selectively and reversibly suppress hippocampal neuronal activity in freely moving animals, allowing excellent temporal control over when the hippocampus is functionally isolated during behavioral experiments (49–51). These temporary inactivation methods have proven particularly valuable because they reveal that the rodent hippocampus is necessary for distinct stages of memory processing, whereas permanent lesion studies often show spared function due to compensation by other brain structures (52). Inhibitory interneurons are promising candidates for controlling the output of the hippocampus (53), however, due to the convoluted 3D structure of this brain region, effective viral vector delivery without spillover and subsequent experimental contamination from neighboring regions challenging. The *hippo BASK-IT* approach should help overcome this limitation.

Future studies should first carefully evaluate the potential off-target consequences of the sparse amygdala labeling. Refinement of the technique through AAV particle titration may further reduce off-target expression. Further examination of whether specific basket cell subclasses are targeted is also an important next step. Chemogenetic variants of this vector could provide not only a significant advancement in basic science towards understanding neuronal circuits, but also a possible opportunity for establishing novel therapeutic methodologies for treating pathophysiological conditions involving hippocampal hyperexcitability, such as epilepsy and Alzheimer’s disease.

## Materials and Methods

Young adult (6–12 week old) C57Bl/6J mice of both sexes were used for experiments, with data collected from ≥3 mice per experimental condition. C57Bl/6J mice were originally purchased from The Jackson Laboratory (RRID:IMSR_JAX:000664) and breeding colonies are maintained in our Division of Animal Resources facility at Emory. All animal procedures were approved by the Emory University Institutional Animal Care and Use Committee (IACUC).

### Enhancer-AAV

The enhancer (AiE0476m) was identified using the Allen Institute brain knowledge platform:

GAGAAACACCACAAGCAAAGCCAAAATAGGTGGCCTAGAACTTCCAGCTTGAAATATGGGAGAGAATGAGGGAGGCACTGTAGAGCAGCTGCCGGGTGCCGCATGAGAACAATTCTCCCTGCTCATAATTAATCCTACCTATTTCTGATGACAGCTGGCTCTTCACTTTGAACAAGCTAGTTAACAACTTTCTTCTCACATTGAGCAAATAATTCATATTTAATTACTTAACCACCAGTTACAAAATGAGAATCATCAAGGA ATCACAATTAATTTGCTATTGACAAACTCATACTTTTAGCAGGCTGATTTCTACTTTATACTTAGATTGGTAATGAAAAATGAAGCTTATTTTAGTTGATTGGTTGGACTTGTGTATGAATATTATCTATTATTTGAAAAGCCAAACTTGAATGCAAAAAAATATTGAATATGAAAA

The enhancer-AAV expressing TurboFP635 (also known as Katushka) was packaged into the PHP.eB AAV serotype. Two copies of TurboFP635 were placed in tandem to increase brightness, with the downstream ORF being codon-optimized to differentiate from the first.

### Retro-orbital injections

Mice were given PHP.eB AAV intravenous injections through the retro-orbital route which crosses the blood-brain barrier as previously described in Chan et al., 2017 (25). Mice were first anesthetized with 2.0% isoflurane (~1 min) to complete unconsciousness to reduce stress during the injection. The vector was titrated to approximately 1.0 × 10^11^ viral genome copies in sterile saline (final volume 65 μl/mouse). Injections of the titrated virus were performed in the left retro-orbital sinus using a 31G × 5/16 TW needle on a 3/10 mL insulin syringe. Mice were kept on a heating pad for the duration of the procedure until recovery (~1 min), followed by a 4-week viral expression period until sample collection.

### Slice preparation and tissue fixation

For immunohistochemistry, mice were fully anesthetized with isoflurane and sacrificed by cardiac perfusion (PBS ~10mL followed by 4% PFA ~ 25mL). The brain was then removed by dissection and placed in 4% PFA solution for 45 minutes. Brain slices were sectioned in phosphate buffer solution (PBS) using a vibrating blade microtome (VT1000S, Leica Biosystems). 50 µm thin coronal slices were cut and subsequently stored in PBS solution. For electrophysiological recordings, mice were fully anesthetized with isoflurane and sacrificed by decapitation. The brain was immediately removed by dissection and submerged in ice-cold cutting solution (in mM) 87 NaCl, 25 NaHCO3, 2.5 KCl, 1.25 NaH2PO4, 2 MgCl2, 1 CaCl2, 10 glucose, and 75 sucrose. Brain slices (100–200 μm) were sectioned using a vibrating blade microtome (VT1200S, Leica Biosystems) in the same solution. Slices were transferred to an incubation chamber and maintained at 34°C for ~30 min and then at 23–24°C thereafter. During whole-cell recordings, slices were continuously perfused with (in mM) 128 NaCl, 26.2 NaHCO_3_, 2.5 KCl, 1 NaH_2_PO_4_, 1.5 CaCl_2_, 1.5 MgCl_2,_ and 11 glucose, maintained at 30.0°C ± 0.5°C. All solutions were equilibrated and maintained with carbogen gas (95% O_2_/5% CO_2_) throughout.

### Immunohistochemistry

Free-floating sections were incubated in 0.1% (v/v) Triton X-100 and 3% (w/v) hydrogen peroxide to eliminate endogenous peroxidase, rinsed in PBS, and pre-blocked in 5% (v/v) normal donkey serum for 30 min at room temperature (RT). In a subset of the sections, parvalbumin immunostaining was performed. These sections were permeabilized with PBST and 0.3% Triton-X and pre-blocked by normal goat serum for one hour. Sections were incubated with guinea pig anti-PV antibody (1:2000, 195 308 Synaptic Systems) at room temperature for two hours. Following 4 wash cycles with PBS, goat anti-guinea pig Alexa 488 secondary antibody was administered for two hours. Slices were mounted on microscope slides using Fluoromount-G with DAPI (00-4959-52, Invitrogen). After the recordings, slices were fixed in 0.1 M PBS containing 4% paraformaldehyde at 4°C overnight. Biocytin labeling was visualized with Alexa 488-conjugated streptavidin.

### Electrophysiology

TurboFP635 expressing cells were targeted for somatic whole-cell recording in the CA3 region of the hippocampus by fluorescent lamp illumination (X-Cite 120 PC, Excelitas) and gradient-contrast video microscopy. Electrophysiological recordings were obtained using Multiclamp 700B amplifiers (Molecular Devices). Signals were filtered at 6–10 kHz and sampled at 50 kHz with the Digidata 1440B digitizer (Molecular Devices). For whole-cell recordings, borosilicate patch pipettes (3-5 MΩ tip resistance) were filled with an intracellular solution containing (in mM) 138 potassium gluconate, 2 KCl, 10 HEPES, 4 MgCl_2_, 4 NaATP, 3 L-ascorbic acid, 0.5 NaGTP, and 8 biocytin (pH 7.3; 270–290 mOsm). Pipette capacitance was neutralized in all recordings and electrode series resistance was compensated using bridge balance in current clamp. Liquid junction potentials were uncorrected. Recorded cells received square pulse current injections with 0 pA bias current, starting from −100 pA, with 20 pA increments.

### Imaging and image analysis

Wide-field images were taken using the Nikon Eclipse Ti-E Inverted LED microscope. High-resolution images were collected using the Nikon CSU-W1 SoRa Spinning Disk system in the SoRa4 microlensing configuration. The CSU-W1 system is equipped with four laser lines (385 nm, 475 nm, 594 nm, and 621 nm). Oil immersion 60X (CFI60 PLAN APOCHROMAT LAMBDA D 60X Oil) objective was used for imaging and data was collected using a Hamamatsu ORCA-Fusion Gen III sCMOS camera. To estimate the fraction of virus expressing neurons that were co-labeled by immunohistochemistry against PV, stitched images of the entire hippocampus, including the DG, CA3, CA2 and CA1 were used, as captured by a 10x objective. The mean fluorescence of the hippocampal PV immunostaining was determined, and the images were baseline subtracted using the mean plus three-times standard deviation. Red (TurboFP635+) cell bodies were selected by hand and PV positivity was established using the baseline subtracted images. Cell bodies were collected from the DG, CA3, and CA1. PV+ cell bodies were collected from every layer without discrimination of the hippocampal subfields.

